# A burns and COVID-19 shared stress responding gene network deciphers CD1C-CD141-DCs as the key cellular components in septic prognosis

**DOI:** 10.1101/2023.02.17.528914

**Authors:** Qiao Liang, Lei Wang, Jing Xu, Anqi Lin, Yongzheng Wu, Qing Tao, Bin Zhang, Haiyan Min, Shiyu Song, Qian Gao

## Abstract

Differential body responses to various stresses, infectious or noninfectious, govern clinical outcomes ranging from asymptoma to death. However, the common molecular and cellular nature of the stress responsome across different stimuli is not described. In this study, we compared the expression behaviors between burns and COVID-19 infection by choosing the transcriptome of peripheral blood from related patients as the analytic target since the blood cells reflect the systemic landscape of immune homeostasis. We identified an immune co-stimulator (CD86)-centered network, named stress-response core (SRC), which coordinated multiple immune processes and was robust in membership and highly related to the clinical traits in both burns and COVID-19. An independent whole blood single-cell RNA sequencing of COVID-19 patients demonstrated that the monocyte-dendritic cell (Mono-DC) wing was the major cellular source of the SRC, among which the higher expression of the SRC in the monocyte was associated with the asymptomatic COVID-19 patients, while the quantity-restricted and function-defected CD1C-CD141-DCs were recognized as the key signature which linked to bad consequences in COVID-19. Specifically, the proportion of the CD1C-CD141-DCs and their SRC expression levels were step-wise reduced along with worse clinic conditions while the sub-cluster of CD1C-CD141-DCs of the critical COVID-19 patients was characterized of IFN signaling quiescence, high mitochondrial metabolism and immune-communication inactivation. Thus, our study identified an expression-synchronized and function-focused gene network which was decreased under burns and COVID-19 stress and argued the CD1C-CD141-DC as the prognosis-related cell population which might serve as a new target of diagnosis and therapy.

## Introduction

The stresses can evoke comprehensive and dynamic responses in host upon the severity of damages, times of actions and the nature of stimuli, and result various outcomes ranging from asymptoma to deaths (Yaribeygi, Panahi, Sahraei, Johnston and Sahebkar 2017). However, the underlying mechanisms of stress-responses which dictate the clinic outcomes remain unclear. Clinically, a coupled manifestation characterized as hyper-inflammation and hypo-immune response represents the severest dyshomeostasis of stress responses and often leads to the worst prognosis with aggravated damages to organs and subsequent multiple organ failure (MOF), systemic inflammatory response syndrome (SIRS) or sepsis, suggesting that the disrupted immune homeostasis, presumably involving the alterations of specific cell type(s), their number and molecular dysfunction, might serve as the critical and common step mediating worsen progressions of various stresses (Hosseini, Hashemi, Shomali, Asghari, Gharibi, Akbari, Gholizadeh and Jafari 2020; Lord, Midwinter, Chen, Belli, Brohi, Kovacs, Koenderman, Kubes and Lilford 2014; Osuka, Ogura, Ueyama, Shimazu and Lederer 2014; Tsirigotis, Chondropoulos, Gkirkas, Meletiadis and Dimopoulou 2016).

The monocytes/DCs in blood serve as an essential linkage node of innate and adaptive immune responses should be the first suspect (Murray 2018; Qian and Cao 2018; Sun Nguyen and Gommerman 2020). Monocytes exhibit a high degree of plasticity and heterogeneity and play a key role in entire stress responses which functionally differentiate to DCs and macrophages (Yang, Zhang, Yu, Yang and Wang 2014). DCs, as the professional antigen-presenting cells (APCs), are essential in initiation and regulation of the transition of innate to adaptive immune response (Segura 2016; Steinman and Nussenzweig 1980). DCs can be divided into 2 major classes, named plasmacytoid dendritic cell (pDC) and conventional dendritic cell (cDC) (Collin and Bigley 2018). With the development of single-cell sequencing, more precisely classified sub populations of DCs were described including CLEC9A+ DCs, CD1C+_A and _B DCs, CD1C-CD141-DCs, AXL+SIGLEC6+ DCs (AS DCs) and pDCs, while CD1C-CD141-DCs were enriched for type I interferon signaling pathway and anti-virus (Villani, Satija, Reynolds, Sarkizova, Shekhar, Fletcher, Griesbeck, Butler, Zheng and Lazo et al. 2017). It is known that the type I IFN produced from DCs facilitate both virus removal by macrophage and DCs integrity to initiate adaptive immune response (Teijaro, Ng, Lee, Sullivan, Sheehan, Welch, Schreiber, de la Torre and Oldstone 2013; Wang, Swiecki, Cella, Alber, Schreiber, Gilfillan and Colonna 2012; Wilson, Yamada, Elsaesser, Herskovitz, Deng, Cheng, Aronow, Karp and Brooks 2013). The studies also showed that in sepsis, the effective drug protection from death required functional T cells (Chang, Svabek, Vazquez-Guillamet, Sato, Rasche, Wilson, Robbins, Ulbrandt, Suzich and Green et al. 2014; Dirks, Egli, Sester, Sester and Hirsch 2013). These studies suggested that the innate-adaptive immune transition and homeostasis may be the common key event in stress response essential for life and death. Although various independent studies have reported different details of stress-induced immune disorders, such as imbalances of pro- vs anti-inflammatory, and innate vs adaptive immune responses, the development of high-throughput sequencing and bioinformatics have given new opportunity to detail the cellular and molecular cores under defined stresses and address the shared mechanism of different stimuli.

Burns are mainly characterized by serious destruction of the skin and related deeper tissues, and are often complicated by MOF, SIRS and sepsis, which leads to nearly 180,000 deaths each year (Alipour Mehdipour and Karimi 2020; Jeschke, van Baar, Choudhry, Chung, Gibran and Logsetty 2020). Noted of, the immune dysregulation is a common feature of severe burns which not only occurs in affected tissues but also system (Korzeniowski, Mertowska, Mertowski, Podgajna, Grywalska, Strużyna and Torres 2022; Zhou, Xu, Herndon, Tompkins, Davis, Xiao, Wong, Toner, Warren and Schoenfeld et al. 2010). The damage-associated molecular patterns (DAMPs) or alarmins, as well as extracellular nuclear debris, such as damaged DNA and RNA molecules generated by burn-damaged tissue will activate immune cells (monocytes, DCs, macrophages etc.) within a few hours (Greenhalgh 2019; Nielson, Duethman, Howard, Moncure and Wood 2017; Osuka, Ogura, Ueyama, Shimazu and Lederer 2014). The destructed first line of immune defense (skin, mucosa and various immune cells) makes organism more susceptible to exogenous infection. Pathogen-associated molecular patterns (PAMPs), as well as DAMPs, are further recognized by pattern recognition receptors (PRRs) which activate the downstream inflammatory pathways and release various inflammatory mediators, and, more importantly, trigger innate to adaptive immune progress, including antigen presentation and T cell proliferation (Bartok and Hartmann 2020; Hennessy and McKernan 2021; Tan, Sun, Chen and Chen 2018; Zindel and Kubes 2020). Any predisposition in this process that compromises proper immune response will be highly disruptive.

COVID-19 pandemic, caused by the severe acute respiratory syndrome coronavirus 2 (SARS-CoV-2), has led to more than 614 million cases and 6.5 million deaths worldwide (https://coronavirus.jhu.edu/map.html). The obvious heterogeneous outcomes of it with most patients having mildly ill or even asymptomatic while others developing critical complications and deaths suggested the existence of differential stress responses of individuals (Chen, Klein, Garibaldi, Li, Wu, Osevala, Li, Margolick, Pawelec and Leng 2021; Wang, Hu, Hu, Zhu, Liu, Zhang, Wang, Xiang, Cheng and Xiong et al. 2020; Wiersinga, Rhodes, Cheng, Peacock and Prescott 2020). It is documented that the innate immune system recognizes viruses by PRRs, such as Toll-like receptor (TLR) 7 (activated by single-stranded RNA) (Cervantes-Barragan, Zust, Weber, Spiegel, Lang, Akira, Thiel and Ludewig 2007), retinoic acid-inducible gene I (RIG-I) and melanoma differentiation-associated gene 5 (MDA-5) (activated by cytosolic viral RNA) (Zust, Cervantes-Barragan, Habjan, Maier, Neuman, Ziebuhr, Szretter, Baker, Barchet and Diamond et al. 2011) etc. The downstream of PRRs cascades the expression of pro-inflammatory factors, among which the type I interferons (IFN-α and IFN-β) play a major role in anti-virus (Schneider Chevillotte and Rice 2014). The delay or faultiness of IFNs function leads to uncontrolled viral replication and massive host cell deaths (Prompetchara Ketloy and Palaga 2020; Yang, Du J, Chen, Zhao, Yang, Su, Cheng and Tang 2014). Also, the adaptive immune system provides an another mechanism to clearance viruses by establishing immunological memory. Both CD4+ and CD8+ T cells are responsible for SARS-CoV-2 clearance by MHC-I and MHC-II, respectively (Channappanavar Zhao and Perlman 2014; Chen, Lau, Lamirande, Paddock, Bartlett, Zaki and Subbarao 2010; Koutsakos Nguyen and Kedzierska 2019; Zhao Zhao and Perlman 2010), while antibody produced by B cells is enhanced by CD4+ T cell and further promote CD8+ T-cell-mediated cytotoxicity (Koutsakos Nguyen and Kedzierska 2019). Thus, the disconnection of innate and adaptive immune response, characterized by sustained and amplified activation of innate immune and insufficient adaptive immune response, results in worse outcomes. In fact, reduced number and altered activation of DCs were reported in severe COVID-19 patients with shifting from IFN-I to pro-inflammatory factor cocktail production (Thevarajan, Nguyen, Koutsakos, Druce, Caly, van de Sandt, Jia, Nicholson, Catton and Cowie et al. 2020; Yang, Chu, Hou, Chai, Shuai, Lee, Zhang, Wang, Hu and Huang et al. 2020; Zhou, To, Wong, Liu, Zhou, Li, Huang, Mo, Luk and Lau et al. 2020). The similar responding pattern was as well demonstrated in aging individuals after the stress challenges (Agrawal and Gupta 2011; Wong Magnusson and Ho 2010). The co-stimulatory molecules CD80 and CD86 on cDCs were also compromised which were urgent to T cell proliferation and activation (Zhou, To, Wong, Liu, Zhou, Li, Huang, Mo, Luk and Lau et al. 2020). The recovery of DCs number and function by targeting these molecules is a promising mean to reduce mortality in COVID-19 related sepsis (Borges Hohmann and Borghi 2021).

Herein, we hypothesized that there might be a shared molecular pattern with designated cellular dysfunctions between burns and COVID-19 infection, which is responsible for worse clinic outcomes. To verify this hypothesis, we applied weighted gene co-expression network analysis (WGCNA) and identified a gene module (named skyblue) related to burns initiated stress, progression and outcomes in GSE19743 dataset and further validated the robustness of the gene network and module-traits correlation in another burns dataset GSE182616. Interestingly, the skyblue core genes, referred to stress-response core (SRC), were conserved in membership and specifically suppressed in ICU COVID-19 patients. The SRC genes were mainly expressed in monocytes and DCs upon single-cell RNA sequencing analysis, and among which the CD1C-CD141-DCs exhibited a relationship with COVID-19 severity and outcome. The sub cluster of these DCs from critical COVID-19 patients demonstrated a “non-infected” molecular profile of compromised IFN signaling and was inactive in cell-cell interaction with other immune components upon silico analysis.

## Materials and Methods

### 1. Data Collection and Preprocessing

The peripheral blood transcriptome expression profiles and clinical traits of burns patients, including GSE19743 and GSE182616, were downloaded from GEO database (https://www.ncbi.nlm.nih.gov/geo) by R package GEOquery (Version 2.62.2) (Davis and Meltzer 2007). The transcriptome expression data was quantile-normalized for further analysis. The peripheral blood leukocytes RNA-sequencing of patients with or without COVID-19 and related clinical information (GSE157103) were downloaded from GEO database. The Single-cell RNA sequencing data of COVID-19 patients and healthy control was downloaded from CELLxGENE website (https://cellxgene.cziscience.com/collections/ddfad306-714d-4cc0-9985-d9072820c530).

### 2. WGCNA

WGCNA (Version 1.71) was performed to identify crucial gene co-expression modules related to clinical traits on GSE19743 (Langfelder and Horvath 2008). First, the function “goodSamplesGenes” in R package WGCNA was used to confirm the input genes and samples were suitable, and none outliers were identified. Secondly, Pearson’s correlation analysis of all pairs of genes was used to construct an adjacency matrix. After that, the adjacency matrix was used to build a scale-free network based on a soft-thresholding parameter β which is 8 in the current case, which enhanced strong correlations between genes and penalized weak correlations. Then the matrix was turned into a topological overlap matrix (TOM) to measure the network connectivity of a gene, which was defined as the sum of its adjacency with all other genes. To classify genes with similar expression patterns into gene modules, the average linkage hierarchical clustering was conducted according to the TOM-based dissimilarity measure with a minimum size (gene group) of 30 for the genes dendrogram and 0.1 for mergeCutHeight. The non-co-expressed genes were included in the “gray” module. The relationships of co-expressed gene modules with clinical traits, such as TBSA, survival status and time point, were analyzed.

### 3. Differential Gene Correlation Analysis (DGCA)

R package DGCA (Version 1.0.2) was employed to detect module robustness by comparison the differences in the correlations of gene pairs between distinct biological conditions. Briefly, DGCA calculates the gene pair correlation in each condition and normalizes correlation coefficients by Fisher z-transformation (McKenzie, Katsyv, Song, Wang and Zhang 2016). The finial gene pair correlation status in each condition is determined by empirical p-values computed via permutation testing. Each gene pair is classed as positive (+), negative (−) and non-significant (0) and grouped by combination of specific condition. The function “ddcorAll” was applied in this section.

### 4. Single Sample Gene Set Enrichment Analysis (ssGSEA)

The ssGSEA was carried by R package GSVA (Version 1.42.0) with the function gsva in which the parameter method was set to ‘ssgsea’ (Hänzelmann Castelo and Guinney 2013).

### 5. DEGs and enrichment analysis

R package limma (Version 3.50.3) was applied to identify DEGs with the threshold of |logFC| >1 and p-value <0.05 (Ritchie, Phipson, Wu, Hu, Law, Shi and Smyth 2015). The module gene enrichment analysis was carried out by the R package clusterProfiler (Version 4.2.2) (Yu, Wang, Han and He 2012). Gene Ontology (GO), including biological process (BP), cellular components (CC) and molecular function (MF), and Kyoto Encyclopedia of Genes and Genomes (KEGG) were analyzed. The p-value <0.05 was considered as a significant outcome.

### 6. Transcription factors analysis in bulk transcriptome

R package RcisTarget (Version 1.14.0) was employed to predict transcription factors of skybklue core genes based on hg19-tss-centered-10kb-7species.mc9nr.feather gene-motif rankings database (Aibar, Gonzalez-Blas, Moerman, Huynh-Thu, Imrichova, Hulselmans, Rambow, Marine, Geurts and Aerts et al. 2017). Motifs were annotated to TFs based on the pathway enrichment analysis, and the highest normalized enrichment score (NES) was picked up for each TFs. The transcription regulation network was visualized by gephi software (Version 0.9.7).

### 7. Single-cell analysis

#### Single-cell sequencing data download, preprocessing and cell annotation

The COVID-19 single-cell sequencing data was obtained from CELLxGENE website. The cells from COVID-19 patients and healthy control were selected for further analysis. Cells with fewer 200 genes expression and >10% mitochondrial reads were removed in quality control, Then, the SC-RNAseq data was normalized, scaled and dimension-reduction (PCA and UMAP). We set dim=15 to in “FindNeighbors” function to construct nearest-neighbor graph according to the standard deviations of the principle components which was visualized by “ElbowPlot” function. The cells were divided into 15 populations by “FindClusters” function with resolution=0.2. RunUMAP was applied for visualization. The cell type was determined by the cluster marker majorly referred to CellMarker database (http://xteam.xbio.top/CellMarker/) as follows, CD4+ T cell (CD3D, CD3E, CD14, LTB, IL7R, MAL), CD8+ T cell (CD3D, CD8A, CD8B, GZMK, NKG7), NK (GNLY, GZMB, NKG7, KLRD1, KLRF1, CD247), B cell (CD79A, CD74, MS4A1, CD19, HLA-DRA, HLA-DQA1), plasma cell (JCHAIN, IGHG1, IGHA1, MZB1, TXNDC5), monocyte (CD14, CD68, S100A12, SERPINA1), pDC (PLD4, ITM2C, LILRA4, IRF7, IRF8, SERPINF1), cDC (CD1C, CD86, HLA-DRA, HLA-DRB5, HLA-DRB1), CD1C-CD141-DC (MS4A7, CFD, LRRC25, FCGR3A, CD68), HSC (CD34, SPINK2, GATA2, MYB, ALDH1A1), RBC (HBB, HBA2, HBA1, HBM), platelet (CD151, GP9, ITGA2B). R package Seurat (Version 4.1.1) was applied in this section (Hao, Hao, Andersen-Nissen, Mauck, Zheng, Butler, Lee, Wilk, Darby and Zager et al. 2021).

#### SRC genes cell location

The cell-location of SRC genes was determined by expression level and Fisher’s Exact Test (FET) (Song, Agrawal, Von Itter, Fontanals-Cirera, Wang, Zhou, Mahal, Hernando and Zhang 2021). For FET part, we first identified a list of cells expressing the gene with a threshold count_i > 0 for a given gene i in SRC, then tested how these cells were enriched for the cells in each inferred cell type under the comparison of one-to-rest by FET. Adjusted p value by FDR < 0.05 was considered significance to designate a cell type for the gene. The enrichment score was calculated by −log10(p-adjust). And of nation, if the adjusted p value of some cell types which equaled to 0, then the enrichment score of these cell types was set to 1 while others was 0. R package stats (Version 4.2.1) function “fisher.test” was utilized in this part.

#### Single-cell score calculation

The single-cell score was calculated by Seurat function “AddModuleScore” of gene lists (Hao, Hao, Andersen-Nissen, Mauck, Zheng, Butler, Lee, Wilk, Darby and Zager et al. 2021). The search parameter was set to TRUE to match the gene symbols. The gene lists of IFN signaling (IFN alpha/beta and IFN gamma) were downloaded from MSigDB database (http://www.gsea-msigdb.org/gsea/login.jsp) with the accession of RECATOME INTERFERON ALPHA/BETA and RECATOME INTERFERON GAMMA.

#### Fractional abundance of cell population analysis

The confidence intervals of relative proportion of different cell types was calculated by R package REdaS (Version 0.9.4) function “freqCI”.

#### CD1C-CD141-DC sub cluster analysis

This section was achieved by R package Seurat (Hao, Hao, Andersen-Nissen, Mauck, Zheng, Butler, Lee, Wilk, Darby and Zager et al. 2021). First, CD1C-CD141-DCs were filtered and followed by Seurat pipeline analysis, including normalization, scale and dimension-reduction. The sub clusters of CD1C-CD141-DC were decided by function “FindClusters” which the resolution was set to 0.3.

#### Single-cell regulatory network inference and clustering (SCENIC) analysis

The SCENIC analysis was applied by pySCENIC (version 0.11.2) (Aibar, Gonzalez-Blas, Moerman, Huynh-Thu, Imrichova, Hulselmans, Rambow, Marine, Geurts and Aerts et al. 2017). The expression matrix of raw UMI counts of CD1C-CD141-DCs was employed in this section. After default data filtering, grnboost2 method was utilized to generate gene regulatory networks. Then, the enriched motifs were determined by “ctx” function in pySCENIC based on cisTarget Human motif database v9 of regulatory features 10 kb centered on the TSS. Finally, enrichment of regulons across single cells was scored by the “aucell” function. The results were visualized by R package ComplexHeatmap (Version 2.10.0) (Gu Eils and Schlesner 2016).

#### Pseudotime analysis

The CD1C-CD141-DC sub clusters were employed to perform trajectory analysis by R package monocle (Version 2.22.0) (Trapnell, Cacchiarelli, Grimsby, Pokharel, Li, Morse, Lennon, Livak, Mikkelsen and Rinn 2014). The different genes (q value <0.01) among the COVID-19 status were picked up to order cells which were calculated by function “differentialGeneTest” of monocle. The genes correlated with pseudotime under the threshold of Spearman correlation index > 0.4 or < −0.4 were selected for visualization.

#### Cell-cell interaction analysis

The cell-cell interaction of CD1C-CD141-DC sub cluster and other cells were achieved by R package CellChat (Version 1.4.0) based on public repository of ligands, receptors, cofactors, and their interactions (Jin, Guerrero-Juarez, Zhang, Chang, Ramos, Kuan, Myung, Plikus and Nie 2021). Briefly, significant interactions of ligand-receptor pairs between cell types were identified by perturbation testing and further constructed communication network. The different incoming and outgoing signals of among CD1C-CD141-DC clusters were systematically analyzed and the specific contributed signaling pathway visualized through bubble plot.

### 8. Statistical analyses

R (Version 4.1.2) was applied to all statistical tests. Spearman correlation analyses were done by R. The statistical analysis of ssGSEA score in bulk transcriptome between different disease status were achieved by Wilcoxon signed-rank test. The correlation index of different skyblue subset genes was calculated by Pearson correlation test. P values of multiple comparisons were adjusted by “FDR”. The statistical analysis of single-cell score among the different cell types or clusters was determined by Kruskal-Wallis test. All P-values were considered significant if <0.05.

## Results

### 1. Identify a clinically relevant gene co-expression network in burns datasets

We initially set to identify clinical traits related co-expression genes by applying weighted gene co-expression network analysis (WGCNA) in burns peripheral blood cell transcriptome dataset GSE19743. The analysis workflow is provided in Supplemental Fig. S1. A total of 114 burns patients and 63 healthy controls were separated into two clusters under the whole transcriptome dendrogram clustering (Fig. 1A), indicating that burns extremely shifted the gene expression. With a scale-free network based on a soft-thresholding parameter β (which was 8 in this case) and topological overlaps, we generated 52 gene co-expression modules according to the hierarchical clustering tree and automatic module detection (Supplemental Fig. S2A). The modules that were correlated with all of total body surface area (TBSA) injured percentage, survival status and hours post burns (time point) were regarded as the vital signatures, which left three modules, namely brown4, pink and skyblue (Fig. 1B, D). Among the three modules, the eigengene of pink and skyblue was either negatively or positively correlated to TBSA or survival status, respectively, demonstrating that these two modules stood for good prognosis, while brown4 showed the opposite (Fig. 1B, D). It is noteworthy that the skyblue module was exclusively related to stress-related traits, with no association with any other clinical factors such as age or gender (Supplemental Fig. S1).

**Figure 1.**
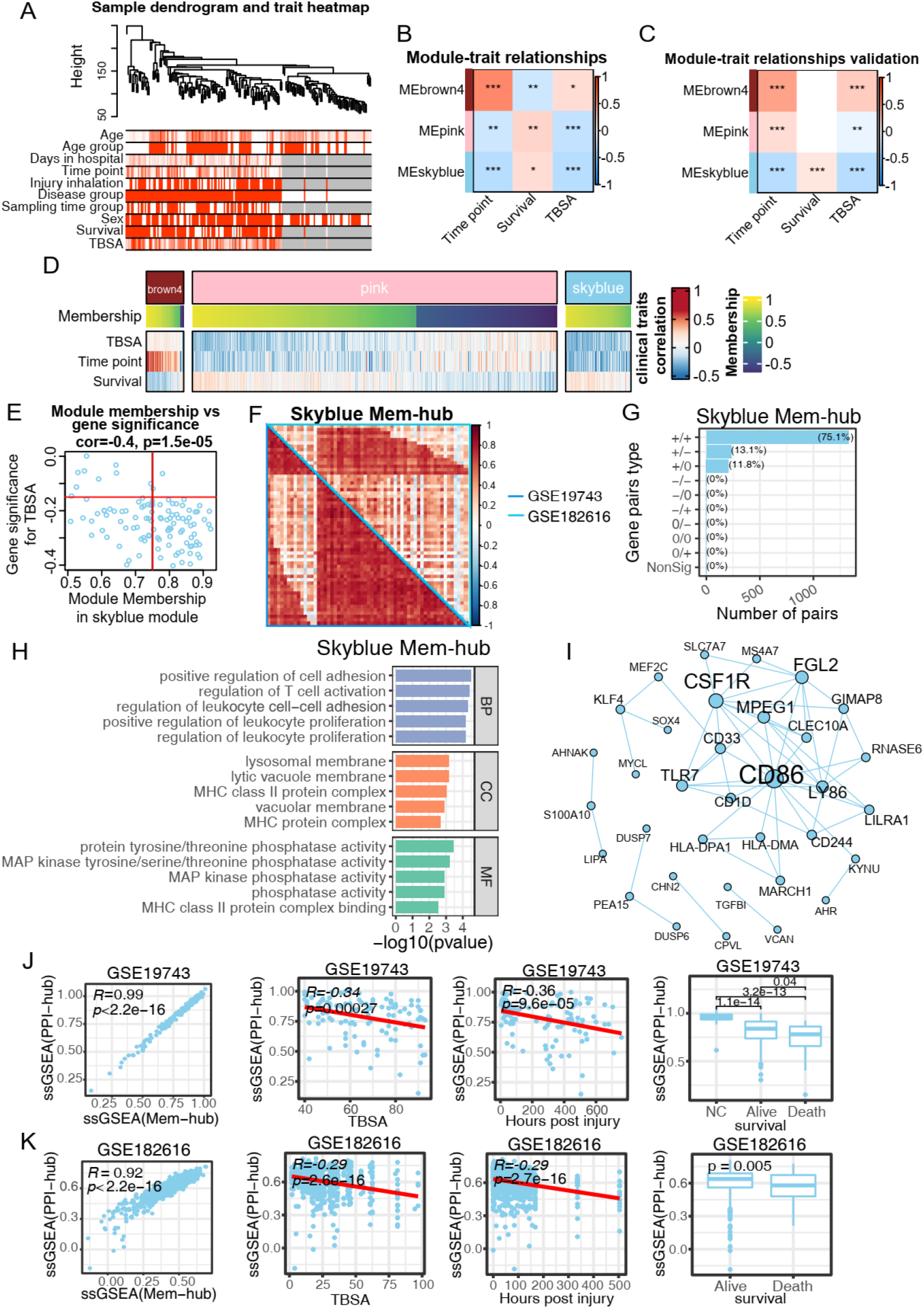
CD86-centered gene network was the robust and prognostic core signature of burns systematic alternation. (A) Clustering dendrogram of samples in burns dataset (GSE19743) based on expression pattern. The clinical information was represented by red color whose intensity was proportional to older age, longer hospital-stay days, later time point of sample-collection, injury inhalation, burns condition, male, survival as well as larger TBSA. (B-C) Representative heatmap of vital module eigengenes and clinical trait correlation in GSE19743 (B) and GSE182616 (C). (D) Heatmap of genes membership and clinical-traits correlation in the vital modules. (E) Skyblue module genes membership and correlation with TBSA. The red line represented the threshold of mem-hub pick-up which was 0.75 and −0.15 for membership and gene significance for TBSA, representatively. (F) Pearson correlation heatmap of skyblue mem-hub genes of GSE19743 and GSE182616 datasets. The left-bottom and right-top part of heatmap is based on GSE19743 and GSE182616, respectively. (G) Barplot of DGCA based on skyblue mem-hub genes between GSE19743 and GSE182616. Each gene pair is classed as positive (+), negative (−) and non-significant (0) and grouped by combination of GSE19743 and GSE182616 datasets. (H) PPI Network of skyblue mem-genes. The size of node and gene label represent the degree of specific node. (I) Barplot of top skyblue mem-hub genes GO enrichment items ordered by p value. (J-K) SsGSEA scores of skyblue PPI-hub genes and mem-hub and correlation (left), as well as correlated with TBSA (middle-left), time point (middle-right) and survival (right) in GSE19743 (J) and GSE182616 (K). Pearson and Spearman correlation was applied for PPI-hub with mem-hub and clinical traits, respectively. * P < 0.05, ** P < 0.01, *** P < 0.001.

To test the above findings, we recruited another burns peripheral blood cell transcriptome dataset (GSE182616). The gene pair correlation in brown4 and skyblue modules between the two datasets illustrated a more conserved pattern than pink module, where 65% (50.1% and 14.9% for conserved positive and negative correlation, respectively) and 65.1% (all for positive correlation) compared to 40.5% (23.3% and 17.2% for conserved positive and negative correlation, respectively) (Supplemental Fig. S2B, C). As for module-trait correlation, only skyblue module stayed firm relationship with all of TBSA, survival status and time post injury (Fig. 1C), claiming that the skyblue module was the robust and vital signature in both module membership and traits correlation.

To condense the skyblue module, we applied membership-traits filtering and protein-protein interaction (PPI) filtering. We first picked up 60 hub genes in skyblue as mem-hub with threshold of membership > 0.75 and gene significance for TBSA < −0.15 (Fig. 1E), which had both higher gene pair correlations and clinical relevance compared to the rest genes. Interestingly, the skyblue mem-hub was more robust between two burn datasets, in which 75.1% gene pairs were consistently positively correlated (Fig. 1F, G), indicating that the mem-hub was functionally central in skyblue. The Gene Ontology (GO) enrichment analysis revealed that the mem-hub was highly involved in immune-related processes, including T cell and leukocyte activation and proliferation, MHC complex formation and MAPK signaling pathway (Fig. 1H), which covered both innate and adaptive immune responses. Remarkably, the mem-hub formatted a condensed PPI network with a few scattered connections (Fig. 1I) according to string website (https://string-db.org/). The network covered multiple immune processes, including pathogen recognition (TLR7), antigen presentation (CD1D, HLA-DMA and HLA-DPA) and adaptive immune co-stimulation (CD86), a bridge molecule of innate and adaptive immune regarded as DCs marker functioned in T cell proliferation and activation and essential for anti-infection (Dyck and Mills 2017; Lal, Rudolph, Dezzutti, Linsley and Prince 1996; Linsley, Brady, Grosmaire, Aruffo, Damle and Ledbetter 1991). The other hub nodes of PPI, such as CLEC10A, CSF1R, MPEG1, FGL2 and LY86, also played essential roles in inflammation and/or immune homeostasis (Hagan, Kane, Grover, Woodworth, Madore, Saleh, Sancho, Liu, Li and Proto et al. 2020; Hou, Wang, Zhou and Li 2021; McCormack, de Armas, Shiratsuchi, Fiorentino, Olsson, Lichtenheld, Morales, Lyapichev, Gonzalez and Strbo et al. 2015; Su, Zhu, Xu, Wang, Dong, Kapuku, Treiber, Gutin, Harshfield and Snieder et al. 2014; Suzuki, Yamamoto, Toyoshima, Osawa and Irimura 1996).

We then extracted 24 genes in sub PPI network (the node number >3) and referred them as PPI-hub. The consistent gene pair correlation of PPI-hub increased from 75.1% to 85.5% compared to mem-hub (Supplemental Fig. S2D), indicating a further centralization of the function of PPI-hub genes. In fact, single sample gene enrichment analysis (ssGSEA) scores of mem-hub and PPI-hub were highly correlated in GSE19743 and GSE182616 whose Pearson correlation index was 0.99 and 0.92, respectively (Fig. 1J, K). The ssGSEA score of PPI-hub was negatively correlated with TBSA and time post injury, suppressed in burns patients and further aggravated in dead ones (Fig. 1J, K). These results implied that PPI-hub was the transcriptional signature related to burns initial injury, progression and prognosis, and functionally enriched in immune response.

### 2. Network genes identified in burns were conserved and prognostic in COVID-19

To test whether there might be similar molecular response to different stresses, we next examined the performance of the “skyblue” PPI-hub from burns in COVID-19 peripheral blood leukocyte transcriptome data. Impressively, the PPI-hub demonstrated a highly synchronous pattern while 96.7% of 276 gene pairs maintained the positive comparable correlation with burns (GSE19743) data (Fig. 2A, B), only nine gene (CD244, CLEC10A, FGL2, HLA-DPA1, KLF4, KYNU, MARCH1, MYCL, TLR7) pairs with RNASE6 were exceptional, emphasized a high similarity of stress response between burns and COVID-19 at gene expression level. Notably, ssGSEA score of PPI-hub was specifically and dramatically restrained in severe COVID-19 patients rather than mild ones, which was supported by both clinical detection and sickness scores (Fig. 2C, D). Specifically, the higher PPI-hub ssGSEA score implied the shorter hospitalizations and less ventilator, while the COVID-19-severity-related information, including acute physiologic assessment and chronic health evaluation (APACHE II) score, sequential organ failure assessment (SOFA) score, and laboratory measurements of C-reactive protein (CRP), D-dimer, ferritin, lactate and procalcitonin claimed a negative correlation with ssGSEA score of PPI-hub (Fig. 2D).

**Figure 2.**
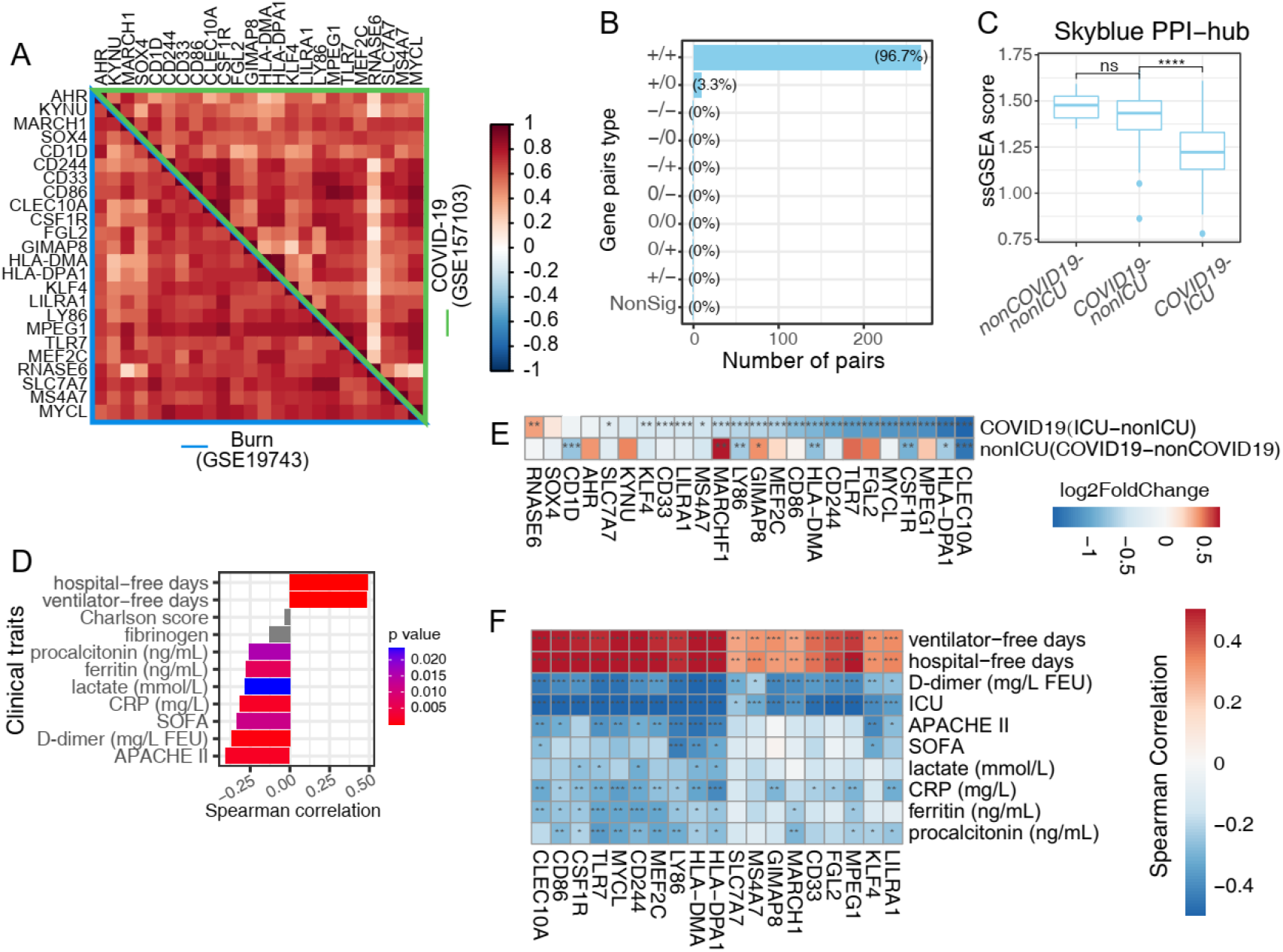
Burns’ core gene net was conserved in both membership and clinical-trait correlation in COVID-19. (A) Pearson correlation heatmap of skyblue PPI-hub genes in burns (GSE19743) and COVID-19 (GSE157103) datasets. The left-bottom and right-top represent burns and COVID-19 datasets and marked by blue and green color, respectively. (B) Barplot of skyblue PPI-hub gene pairs differential correlation on burns and COVID-19 datasets based on DGCA analysis. Each gene pair is classed as positive (+), negative (−) and non-significant (0) and grouped by combination of GSE19743 and GSE157103 datasets (C) Boxplot of ssGSEA sore of skyblue PPI-hub genes across disease conditions in COVID-19 GSE157103 datasets. (D) Barplot of Spearman correlation index of skyblue PPI-hub ssGSEA score with COVID-19 patients clinical traits. The bar color represents the Spearman correlation p values and p ≥ 0.05 was colored by grey. (E) DEG log2FoldChange heatmap of skyblue PPI-hub genes in GSE157103 dataset. The color indicated the log2FoldChange in each disease status comparison. (F) Spearman correlation coefficient heatmap of SRC genes in GSE157103 dataset. * P < 0.05, ** P < 0.01, *** P < 0.001.

We next compared the expression levels of individual PPI-hub genes between different clinic conditions. With no accident, the PPI-hub genes were more widely repressed in ICU COVID-19 patients (19 genes) compared with mild ones in which only 6 genes (CD1D, CLEC10A, CSF1R, HLA-DMA, HLA-DPA1 and LY86) were down-regulated compared with the controls (Fig. 2E). We further simplified PPI-hub by drawing the 19 lower expressed genes in COVID-19 ICU vs mild COVID-19 (AHR, CD1D, KYNU, RNASE6 and SOX4 were struck out), and referred them as stress response core (SRC). Importantly, the individual genes of SRC were also broadly correlated with clinical traits such that the higher expression of which implied the lesser severity of the disease (Fig. 2F). The high correlation coefficients of ssGSEA scores of SRC and PPI-hub gene sets in all of burns and COVID-19 datasets (data not shown) confirmed that SRC was the functional center of the skyblue module.

### 3. COVID-19 scRNA-seq analysis revealed SRC genes monocyte-DC specific

To identify the source cells of SRC genes, we employed single-cell RNA sequencing (scRNA-seq) data of COVID-19 PBMCs. After quality control (QC), a total of 623375 cells from healthy controls, asymptomatic, mild, moderate, severe and critical COVID-19 patients were grouped into 15 clusters by a shared nearest neighbor (SNN) modularity optimization based clustering algorithm (Fig. 3A). The cell clusters were then annotated into 12 cell types according the cluster gene markers (Fig. 3B, Supplemental Fig. S3A). We then examined the expression level and specificity of SRC genes of each cell type in healthy control. 18 of 19 SRC genes were detected in COVID-19 scRNA-seq data except MARCHF1 which was an E3 ubiquitin-protein ligase and constitutively mediated ubiquitination of CD86 and MHC class II proteins and promote their subsequent endocytosis and sorting to lysosomes in immature dendritic cells (Bartee, Mansouri, Hovey, Gouveia and Fruh 2004; De Gassart, Camosseto, Thibodeau, Ceppi, Catalan, Pierre and Gatti 2008). It is not clear why MARCHF1 was not detected in single cell analysis. CD1C-CD141-DCs, cDCs and monocytes, which comprised 3.4%, 1.8% and 11.3% of all cells, respectively, all exhibited a much higher expression level and percentage compared to other cell populations, suggesting that SRC genes were monocyte-DC (Mono-DC) specific (Fig. 3C). FET inspection confirmed this observation (Fig. 3D). Indeed, cDCs, CD1C-CD141-DCs and monocytes had the highest SRC score compared with other cells (Supplemental Fig. S4A).

**Figure 3.**
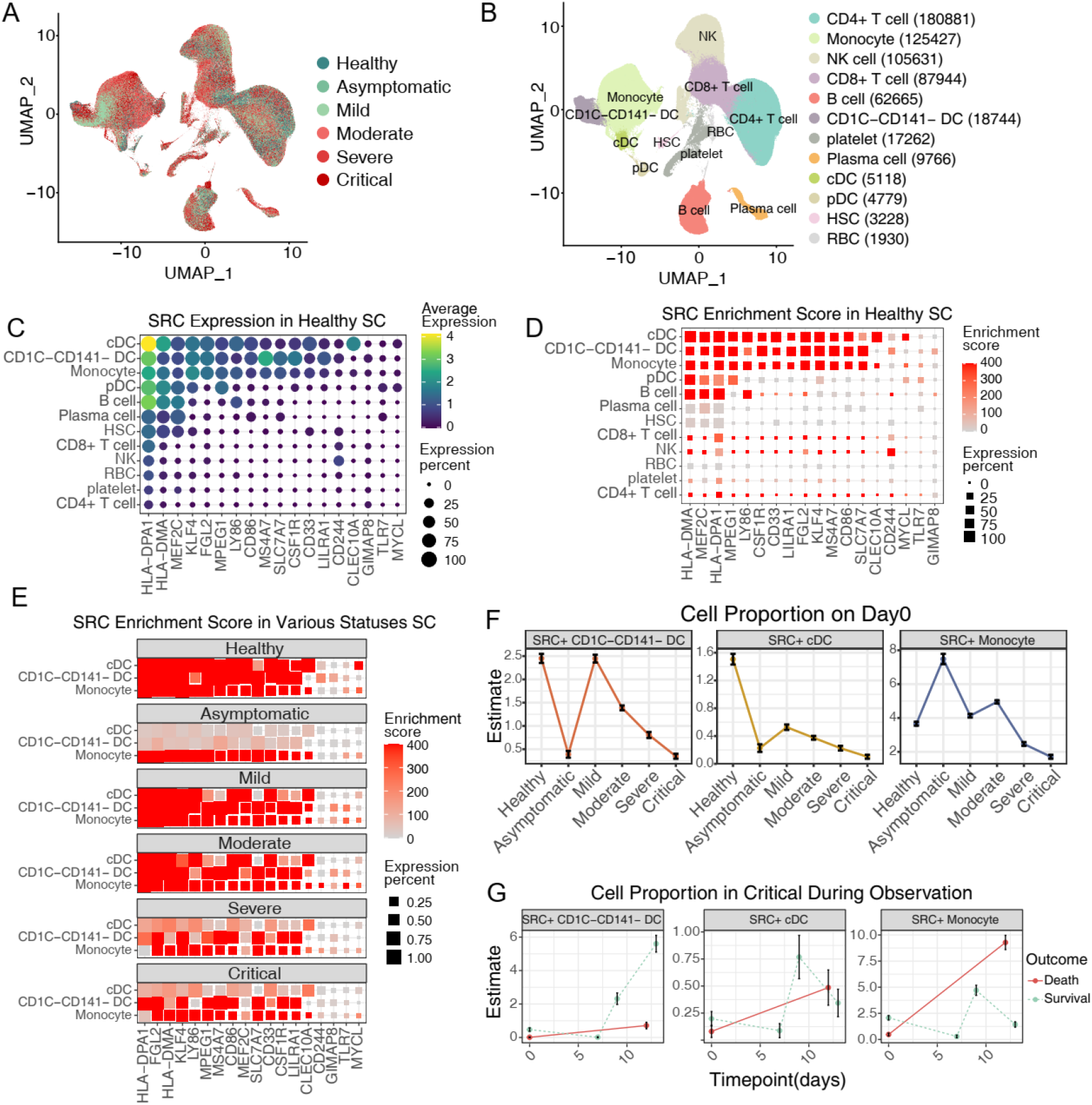
CD1C-CD141-DCs are distinctive and prognostic cell source of SRC genes in COVID-19 scRNA-seq. (A-B) UMAP dimensionality reduction embedding of healthy and COVID-19 patients peripheral blood mononuclear cells after QC and colored by disease status (A) and annotated cell types (B). The numbers indicated the cell number of each cell type in B. (C) SRC genes expression dot plot in healthy subjects. The dot color and size indicated the scaled expression level and percentage. (D) SRC enrichment score of each cell type in healthy subjects. The square color and size represented the enrichment score and expression percentage. (E) SRC enrichment score of CD1C-CD141-DCs, monocytes and cDCs across the disease statuses. The square color and size represented the enrichment score and expression percentage. (F) SRC+ CD1C-CD141-DCs, monocytes and cDCs proportion of different disease statuses on sample-collection time. The error bar represented the 95% confidence interval and the y axis represented the cell percentage. (G) SRC+ CD1C-CD141-DCs, monocytes and cDCs proportion of critical COVID-19 patients during disease progression. The outcome was annotated by line color. The error bar represented the 95% confidence interval. **** P < 0.0001.

We next tested the SRC genes expression pattern across clinical statuses. Remarkably, the SRC genes were broadly expressed by CD1C-CD141-DCs, cDCs and monocytes in healthy controls, mild and moderate COVID-19 statuses while preferred to be expressed by CD1C-CD141-DCs and monocytes in severe and critical COVID-19 statuses (Fig. 3E), indicating that CD1C-CD141-DCs and monocytes were the essential cellular source of SRC in severe and critical COVID-19.

### 4. Enhanced SRC in Mono-DCs showed T-dependent potential and less severity of COVID-19

To assess SRC function in cells, we arbitrarily divided Mono-DCs into SRC positive (SRC+) or negative (SRC-) groups based on the median value of expressed SRC genes (8 as the threshold). Interestingly, both the SRC+ Mono-DCs were functionally enriched in antigen processing and presenting, and T cell activation, suggesting a potential T-dependent function, while SRC- Mono-DCs were enriched in electron transport chain (ETC) and exhibited a tendency to pro-inflammation (Supplemental Fig. S4B-D), indicating that the SRC genes participated the functional polarization of the monocytes.

We next focused on the distributions of SRC+ cells in healthy controls and in various COVID-19 infected patients. We found that the percentages of the SRC+ CD1C-CD141-DCs and cDCs were high, whereas the percentage of the SRC+ monocytes were relatively low in the healthy controls (Fig. 3F). After COVID-19 infection, the percentages of the SRC+ CD1C-CD141-DCs and cDCs exhibited a shifting with a gradual reduction along with the severity of COVID-19 (from mild to critical) (Fig. 3F). Similarly, the proportion of SRC+ monocytes was also high in mild and moderate patients and then gradually reduced along the severity of COVID-19. Thus, these SRC+ Mono-DCs exhibited a stepwise exhausting behavior upon the degree of the infection. Unexpectedly, in asymptomatic COVID-19 individuals, the percentage of the SRC+ monocytes was much higher compared to that in all other clinical statuses (Fig. 3F), whereas the SRC+ CD1C-CD141-DCs and cDCs levels were among the lowest level. Moreover, the antigen presentation score of asymptomatic SRC+ monocytes were comparable to health DCs and higher than that of monocytes from other COVID-19 statuses (Supplemental Fig. S4E). Thus, SRC in monocytes might introduce a specific antigen dependent action and contributed to the COVID-19 asymptomatic phenotype.

It is noticed that the average expression levels of SRC genes in the individual SRC+Mono-DCs showed limited differences across the different clinical statuses (Supplemental Fig. S4F). The SRC scores in overall Mono-DC populations were gradient decreased with COVID-19 severity (Supplemental Fig. S4G), which was consistent with the bulk transcriptome results, proving that it is the reduction of SRC+ Mono-DCs, likely due to the exhaustion of these cells, was vital in COVID-19 progression. To test this hypothesis, we asked whether the recovery of the SRC+ cells proportion was critical for the survival of critical COVID-19 patients. We found that it was the SRC+ CD1C-CD141-DCs (both SRC+ and SRC-) rather than cDCs or monocytes which were dramatically elevated in survived critical COVID-19 patients compared to dead ones (Fig. 3G).

### 5. Poor IFN response and immune-crosstalk of CD1C-CD141-DCs in critical COVID-19

To illustrate the functional features of CD1C-CD141-DCs along with COVID-19 severity, we defined the sub clusters of CD1C-CD141-DCs. With a resolution of 0.3, CD1C-CD141-DCs were subdivided into nine clusters (Fig. 4A). The majority CD1C-CD141-DCs of healthy controls (86.7%) and critical COVID-19 (78.1%) were found in cluster2 and 1, respectively (Fig. 4B), which were juxtaposed in silicon, indicating an overall expression similarity between the cells with no infection or no responsiveness (Fig. 4A). The CD1C-CD141-DCs from other COVID-19 statuses were dispersed in several clusters (Fig. 4A). The cluster1 contained 50.3% cells from critical COVID-19, cluster2 68.4% cells from healthy controls, cluster4 and 6 90.8% and 94.2% cells, respectively, from mild COVID-19 and cluster5 and 8 95.1% and 57.5% cells, respectively, from moderate COVID-19 (Fig. 4B). The critical-COVID-19-dominant cluster1 had the lowest SRC signature score among the clusters (Fig. 4C). Similarly, the status-integrated SRC score gradually reduced with the disease severity and critical COVID-19 had the lowest score (Supplemental Fig. S5E), confirming a close relationship between CD1C-CD141-DCs subclustering, patients’ clinical performance and SRC gene expression.

**Figure 4.**
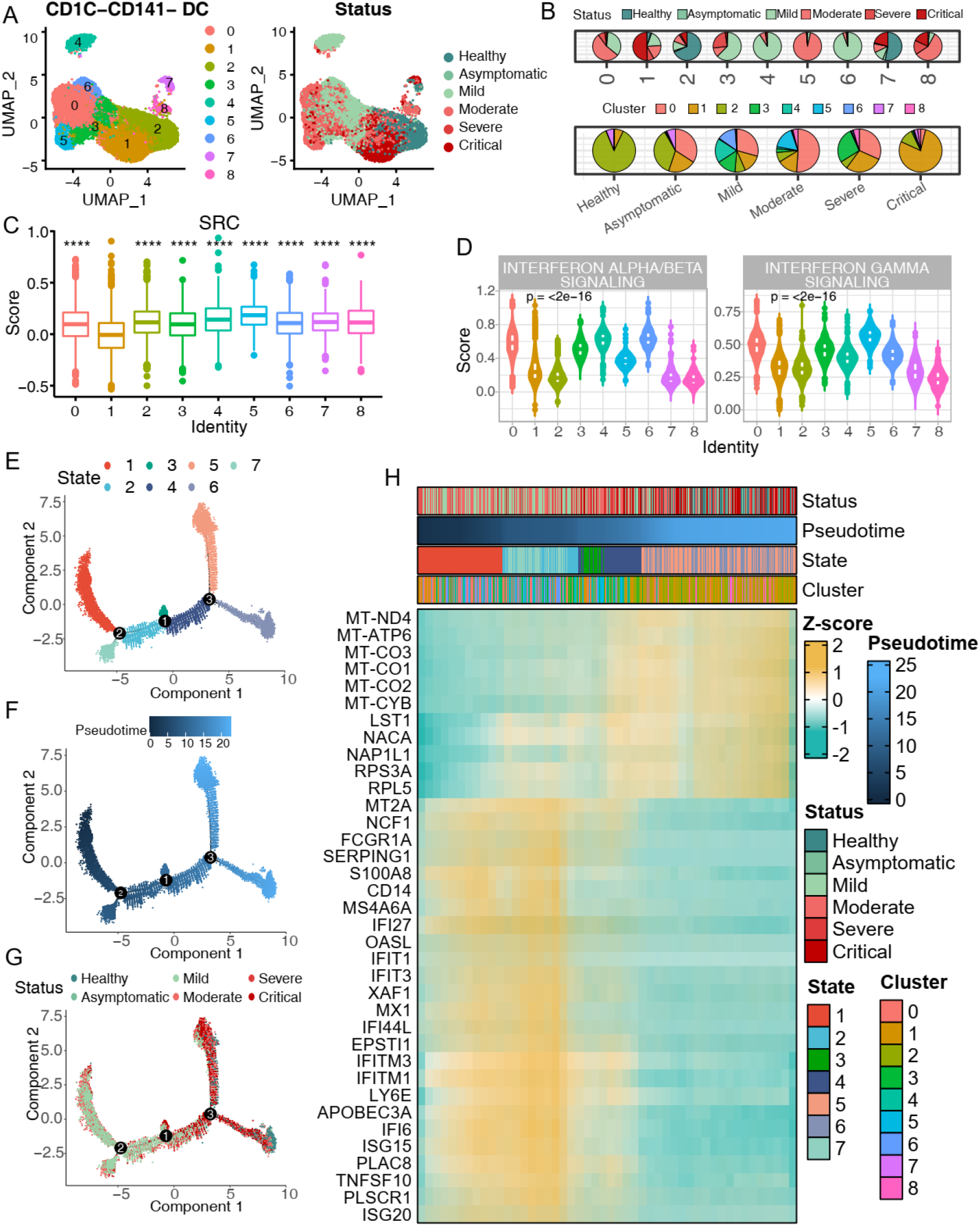
CD1C-CD141-DCs sub cluster of critical COVID-19 patients is IFN-unresponsive. (A) UMAP embedding visualization of CD1C-CD141-DC sub-clusters which colored by cluster identity (left) and disease status (right). (B) Pie chart of CD1C-CD141-DCs composition in each cluster (top) and disease status (bottom), respectively. (C) Boxplot of SRC genes score among each CD1C-CD141-DC sub-clusters. Cluster1 was compared to the other clusters. (D) Violin plot of Interferon signaling score across each CD1C-CD141-DC sub-clusters. Kruskal-Wallis test p value was annotated on the top left. (E-G) UMAP visualization of CD1C-CD141-DC trajectories colored by state (E), pseudotime (F) and disease status (G). (H) Representative heatmap of differentially expressed and correlated with pseudotime genes.

To deduct the inward cellular signaling in CD1C-CD141-DCs, we first applied the transcription factor (TF) activity prediction, which recognized STAT1, SRY, SPI1, SOX9 and IRF8 as the top 5 TFs deducted upon SRC genes (Supplemental Fig. S5A, B). Among these TFs, STAT1 and IRF8 had the highest normalized enrichment score (NES). In scRNA-seq data, we confirmed that the cluster1 CD1C-CD141-DCs was inactivated in STAT1 and IRF8 regulons activity (Supplemental Fig. S5C). ScRNA-seq analysis also showed that the target genes of STAT1 and IRF8 were highly enriched in IFN signaling (both type I and II), antigen presenting and T cell activation, which were down-expressed in both cluster1 (critical) and 2 (healthy) of CD1C-CD141-DCs (Supplemental Fig. S5C, D). Since the IFN signaling pathways were essential for both anti-virus in early infection and regulation of adaptive immune response, we next scored the type I IFN (IFN-α/β) and type II IFN (IFN-γ) signaling pathways in various CD1C-CD141-DCs subclusters. Interestingly in the healthy and critical COVID-19 dominant clusters, both type I and II IFN signaling pathways were inactive (Supplemental Fig. S5E). Moreover, the status-integrated IFN-I and −II signaling scores in critical COVID-19 patients and healthy controls were latent compared to all other COVID-19 statuses (Supplemental Fig. S5E).

For chasing the trajectory of the sub clusters of CD1C-CD141-DCs, we then applied pseudotime analysis by monocle. Consistent with the overall expression pattern dimension reduction by UMAP (Fig. 4A), there was greater transcriptional similarity between CD1C-CD141-DCs from healthy controls and critical COVID-19 patients in trajectory branches and pseudotime (Fig. 4E-G), supporting above observation. Spearman correlation of pseudotime with gene expression revealed that the interferon response genes and mitochondrial genes were preferred to decrease or increase, respectively, in healthy controls and critical COVID-19 ones (Fig. 4H), suggesting these genes as the major contributors in subclustering. Since the suppressed oxidative phosphorylation was the signature of activated DCs (Everts, Amiel, van der Windt, Freitas, Chott, Yarasheski, Pearce and Pearce 2012; Pearce and Everts 2015), this result further supported that the low-responsive or un-stimulated status of the CD1C-CD141-DCs in the critical COVID-19 or healthy controls, respectively. Interestingly, while critical COVID-19 dominant cluster1 was TNF signaling suppressed, including TNFSF10 and TNFSF13B, the healthy control dominant cluster2, as well as all other clusters were not (Supplemental Fig. S4A), highlighting the differences between un-stimulated (health) and non-responsive CD1C-CD141-DCs.

Next, we applied CellChat to elucidate the cell population communications of CD1C-CD141-DC sub clusters, which presumably reflected the role of these cells in regulated stress-response processes of organisms. Overall, the critical COVID-19 dominant cluster1 was most insufficient in both outgoing and incoming signaling count and weight while cluster2 (health) ranked at top among the 9 clusters (Fig. 5A), indicating that the CD1C-CD141-DC sub cluster in critical COVID-19 was immune-communication defective, whereas the CD1C-CD141-DC sub cluster in health was intact. Specifically, the critical COVID-19 dominant cluster1 was disabled in ITGB2, ANNEXIN, ADGRE5, SN and THBS (incoming) and MHC-II, MHC-I, ICAM, TNF, BAFF, SN, GRN and BAG (outgoing) signaling (Fig. 5B, C). Detailed in ligand/receptor pairs, such as in MHC-II signaling, the interaction of HLA-DMB in cluster1 to CD4 in cDCs and monocyte, as well as pDCs was completely missing (Fig. 5C). CD4 signaling in monocyte was previously suggested to promote monocyte to macrophage differentiation and phagocytosis (Zhen, Krutzik, Levin, Kasparian, Zack, Kitchen and Silvestri 2014) while CD4+ monocyte were reduced in COVID-19 (Kazancioglu, Yilmaz, Bastug, Sakalli, Ozbay, Buyuktarakci, Bodur and Yilmaz 2021). Indeed, in our study, we observed that the CD4 expression in monocyte was significantly reduced in severer or critical COVID-19 patients (Supplemental Fig. S6B).

**Figure 5.**
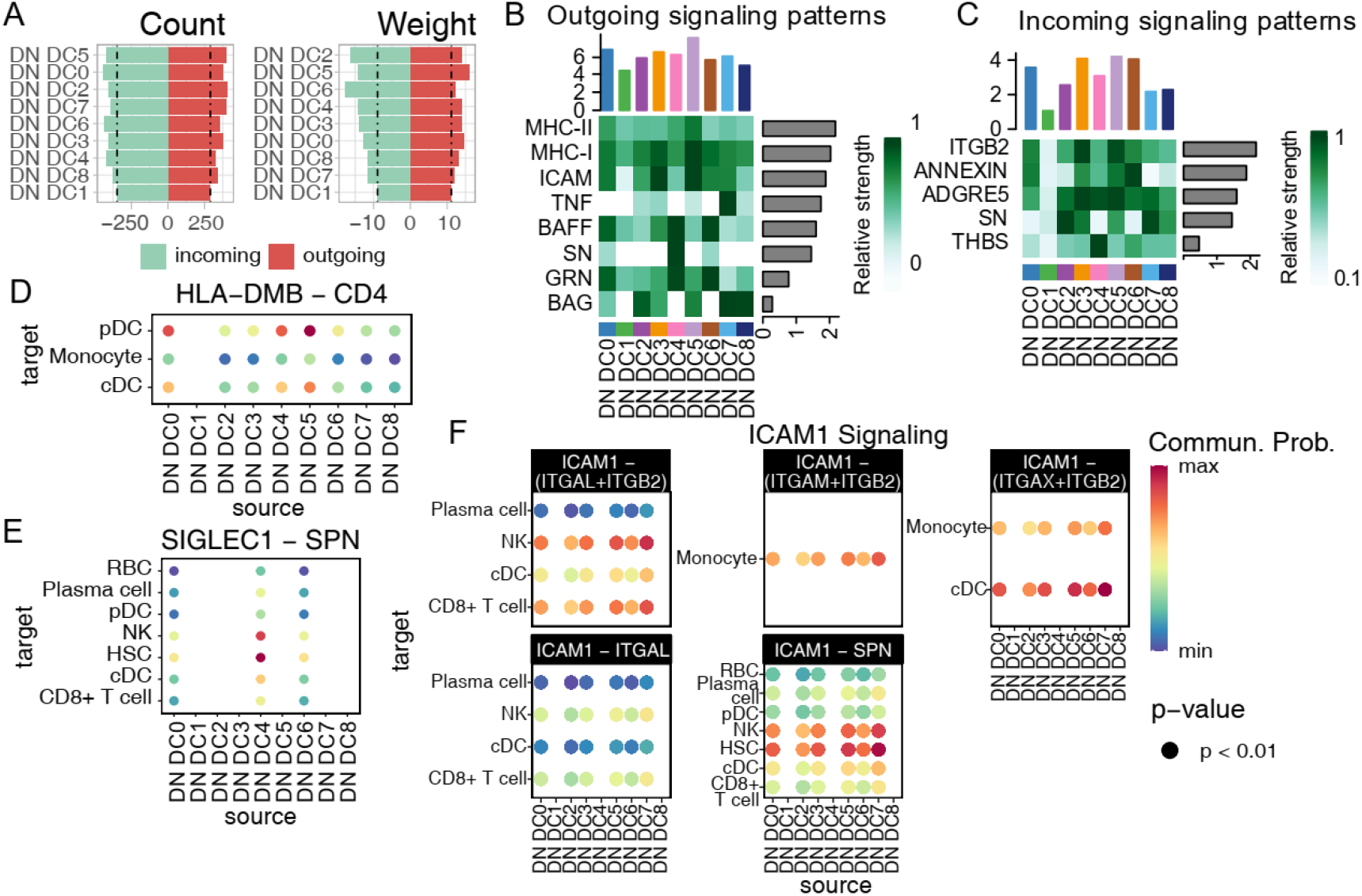
CD1C-CD141-DCs sub cluster of critical COVID-19 patients is immune-crosstalk quiet with other immune cells. (A) Barplot of CD1C-CD141-DC sub clusters overall signaling pathway count (left) and weight (right). The bar was colored by the incoming and outgoing status. DN DC: CD1C-CD141-DC. (B-C) Representative heatmap of CD1C-CD141-DC outgoing (B) and incoming (C) signaling relative strength. (D-F) Representative dot plot of CD1C-CD141-DC outgoing signaling, including HLA-DMB–CD4 (D), SIGLEC1-SPN (E), ICAM1 (F) as well as TNF (G) signaling. The dot size indicated the p value and the color represent communication probability. DN DC represents CD1C-CD141-DC.

In addition, the interaction of SIGLEC1 and SPN (CD43), which was reported to play a vital role in T cell proliferation, differentiation, migration and activation (Galindo-Albarran, Ramirez-Pliego, Labastida-Conde, Melchy-Perez, Liquitaya-Montiel, Esquivel-Guadarrama, Rosas-Salgado, Rosenstein and Santana 2014; Ramirez-Pliego, Escobar-Zarate, Rivera-Martinez, Cervantes-Badillo, Esquivel-Guadarrama, Rosas-Salgado, Rosenstein and Santana 2007), was lost in cluster1 with CD8+ T cells (Fig. 6E). Moreover, the ICAM1 signaling, including interaction with ITGAL, ITGB2 and SPN, was broadly inactive in cluster1 with CD8+ T cells (Fig. 6F). The interaction of ICAM-1 with ITGAL was vital to activate T cells through forming a defined zone in immunological synapse (Grakoui, Bromley, Sumen, Davis, Shaw, Allen and Dustin 1999; Monks, Freiberg, Kupfer, Sciaky and Kupfer 1998). Indeed, CD8+ T cells rather than CD4+ T cells were decreased in severe and critical COVID-19 compared with moderate status (Supplemental Fig. S3C). However, it is worthy to note that the cluser4 (mild COVID-19 dominant) and 8 (moderate COVID-19 dominant) exhibited a similar pattern with cluster1 in ICAM signaling rather than SIGLEC-SPN, indicating the partial dysfunction of cluster 4 and 8 and might be complemented by other clusters. The other signaling pathways were observed different in cluster1, involving TNF, ADGRE5-CD55, ANXA1-FPR2 and THBS1-CD36 (Supplemental Fig. S6C-F), which needed further research.

**Figure 6.**
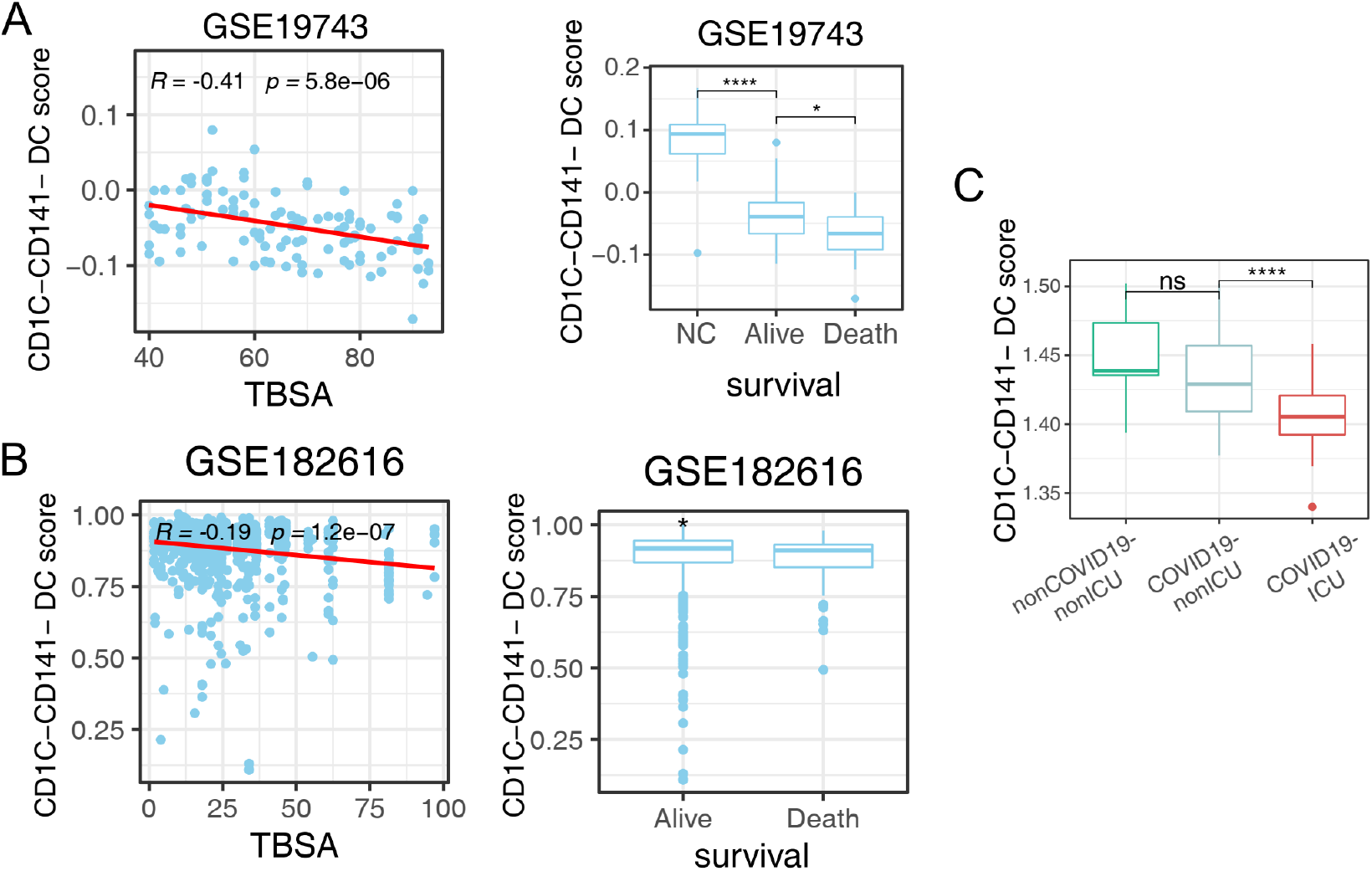
CD1C-CD141-DCs gene markers were related to clinical traits in burn and COVID-19 bulk transcriptomes. (A-B) CD1C-CD141-DCs gene marker scores correlated with TBSA (left) and prognosis (right) in burns transcriptome datasets of GSE19743 (A) and GSE182616 (B). (C) Boxplot of CD1C-CD141-DCs gene marker scores in COVID-19 transcriptome dataset. P values were adjusted by FDR. * P < 0.05, ** P < 0.01, *** P < 0.001.

### 6. CD1-CD141-DC score was related to burns and COVID-19 transcriptome datasets

Finally, to assess whether the single cell results were indeed related to the clinic traits in burns and COVID-19 transcriptome datasets, the CD1-CD141-DC marker genes score was calculated in peripheral blood transcriptome datasets of burns and COVID-19. It was found that the CD1-CD141-DC score was negatively correlated to TBSA in burn datasets and was suppressed in patients with poor prognosis. (Fig. 6A, B). Furthermore, the CD1-CD141-DC score in burn survival patients was dramatically inhibited compared to normal controls, suggesting that the burn has a detrimental effect on CD1-CD141-DCs. Also, the CD1-CD141-DC score shared a similar pattern with the SRC in the COVID-19 transcriptome dataset (Fig. 8A), specifically reduced in critical COVID-19 patients (Fig. 6C). These findings argued that CD1-CD141-DCs were the critical cell population of SRC genes in response to burns and COVID-19.

## Discussion

We hypothesized that different stressors may elicit a shared molecular response. To test this, we identified a CD86-centered network that exhibited coordinated expression and PPI patterns in the transcriptome of the peripheral blood from burn-injured patients using WGCNA. The network included multiple processes related to the innate and adaptive immune response, such as pattern recognition, antigen presentation, and T cell co-stimulation. Remarkably, the genes in the network were robust in membership of expression pattern, and relationship with clinical traits in COVID-19 patients, supporting our initial hypothesis at least in the context of burns and COVID-19 and they were eventually named as SRC (stand for stress-response core).

It is known that the incidence of sepsis in burn patients, typically resulting from infection, is higher (8%–42.5%) than trauma (2.4%–16.9%) or even critical care patients (19%–38%) (Thompson, Zuniga, Sousse, Christy and Gurney 2022). Burn injuries can also increase a patient's susceptibility to infection by compromising their metabolic and immunological defenses (Jeschke, van Baar, Choudhry, Chung, Gibran and Logsetty 2020; Mulder, Koenen, Vlig, Joosten, de Vries and Boekema 2022). Also, burn injuries can expose large amount of nuclear materials, which might initiate a viral-like cellular response. At least, our results demonstrated that multiple immune processes (pattern recognition, antigen presentation, co-stimulation, etc.) were disrupted across the burn initiation, progression and outcome. TLR7 was down expressed in burn murine model whose genetic mutation was observed in severe COVID-19 patients (Butler-Laporte, Povysil, Kosmicki, Cirulli, Drivas, Furini, Saad, Schmidt, Olszewski and Korotko et al. 2022; Wen, Mobli, Radhakrishnan and Radhakrishnan 2022).

The analysis of COVID-19 scRNA-seq demonstrated that the Mono-DCs in the circulation were the cellular host of SRC genes. Among these cells, the CD1C-CD141-DCs were molecularly characterized of enhanced IFN-I and anti-virus compared to other DC populations (Villani, Satija, Reynolds, Sarkizova, Shekhar, Fletcher, Griesbeck, Butler, Zheng and Lazo et al. 2017). Interestingly, CD1C-CD141-DCs were reported to be the most sensitive to exercise-induced mobilization which increased up to 167% after exercise (Brown, Campbell, Wadley, Fisher, Aldred and Turner 2018), indicating that CD1C-CD141-DCs were potentially the fast responders and effectors during stress. Our study also found that CD1C-CD141-DCs deficiency was linked to progression and poor prognosis of COVID-19. Unexpectedly, in the asymptomatic patients, it was the monocytes that expressed higher levels of SRC genes were significantly enriched, while CD1C-CD141-DCs or cDCs were greatly reduced. However, the casual inference of CD1C-CD141-DCs with COVID-19 prognosis, as well as monocytes in asymptomatic patients of COVID-19, and the specific mechanism of mobilization of CD1C-CD141-DCs and monocytes in the disease needed further investigations.

Besides the quantity restriction, the function of CD1-CD141-DCs in critical COVID-19 patients was “poisoned” which was captured as poor IFN response, high mitochondrial metabolism and immune cross talk quiescent. Both type I and II IFNs were effective to defense virus and regulate immune response yet an uncontrolled IFNs led to hyper-inflammation. The impaired IFN signaling, including delayed responses and genetic mutations, was essential for virus replication and tissue damages in COVID-19 (Aso, Ito, Koyanagi and Sato 2019; Yang, Wang, Hui, Yarovinsky, Badeti, Pham and Liu 2021; Zhang Zhao and Zhao 2021). There were 13 subtypes of IFN-α binding to the same receptor which induced different interferon-stimulated genes (ISGs) to modulate antiviral and inflammatory effects (Yang, Wang, Hui, Yarovinsky, Badeti, Pham and Liu 2021). Moreover, the sensitivity and ISGs expression prolife of IFN varied among different cell types (Aso, Ito, Koyanagi and Sato 2019), indicating that the cell specificity to IFN played a key role in IFN double-edge effects (immune regulation vs. hyper-inflammation). DCs, stimulated by IFNs, orchestrated the innate and adaptive immune responses which transformed to the phenotype of high antigen presentation and co-stimulation (Duong, Fessenden, Lutz, Dinter, Yim, Blatt, Bhutkar, Wittrup and Spranger 2022; Lapenta Gabriele and Santini 2020a; Lapenta Gabriele and Santini 2020b). Our studies confirmed the above molecular pattern and further recognized CD1C-CD141-DCs as the vital subtype among the different Mono-DC types in COVID-19 progression and prognosis. Mitochondrial metabolism, including ETC and oxidative phosphorylation (OXPHOS), was suppressed in monocytes/DCs during their activation (Pearce and Everts 2015). CD1C-CD141-DCs in critical COVID-19 and healthy controls both highly expressed ETC components, indicating their unresponsiveness and un-stimulatory status, respectively. Given the fact that the CD1C-CD141-DCs from critical COVID-19 was largely limited to cluster1 which were characterized as immune-crosstalk quiescent, we suggested that the dysfunction of CD1C-CD141-DCs played a key role in critical COVID-19 development and prognosis, though the multi-cell communication patterns in various clusters were complicated likely involving combinatory mechanisms functionally, more studies are guaranteed. However, due to the limitation of burns scRNA-seq data, we are not able to validate the related results in burns condition at cellular level though the CD1-CD141-DC score shared a similar pattern with the SRC genes in the burns and COVID-19 transcriptome datasets, which requires further studies.

## Author Contributions

QL, SS, QG designed the study and wrote the manuscript. QL and SS ran the bioinformatics analyses. LW, JX, AL, QT, BZ, YW and HM provided clinical specimens and data, technical support and conceptual advice.

## Data Availability Statement

Publicly available datasets were analyzed in this study. The transcriptome datasets of burns and COVID-19 can be found in GEO database (https://www.ncbi.nlm.nih.gov/geo/) through the accession numbers of GSE19743, GSE182616 and GSE157103. The scRNA-seq data of COVID-19 patients and healthy control was downloaded from CELLxGENE website (https://cellxgene.cziscience.com/collections/ddfad306-714d-4cc0-9985-d9072820c530).

## Ethics Statements

No animal or human studies are presented in this manuscript.

## Acknowledgements

We thank Nanjing Lupine (YuShanDou) Biomedical Research Institute Co. Ltd., and all the members for their scientific support and bioinformatics analysis advice. We thank Dr. Jianming Zeng (University of Macau), and all the members of his bioinformatics team, biotrainee, for generously sharing their experience and codes.

**Figure Legends**

**Figure S1. Workflow of investigation.** SRC, stress-response core.

**Supplemental Figure S1. Workflow of investigation.** SRC, stress-response core.

**Supplemental Figure S2. Identification and validation of robust gene co-expression module among burns datasets.**

(A) Spearman correlation heatmap of module eigengenes with clinical traits in burns dataset GSE19743.

(B) Pearson correlation heatmap of skyblue module genes of GSE19743 and GSE182616 datasets. The left-bottom and right-top part of heatmap is based on GSE19743 and GSE182616, respectively.

(C) Barplot of DGCA based on skyblue module genes between GSE19743 and GSE182616. Each gene pair is classed as positive (+), negative (−) and non-significant (0) and grouped by combination of GSE19743 and GSE182616 datasets.

(D) Barplot of DGCA based on skyblue PPI-hub genes between GSE19743 and GSE182616. Each gene pair is classed as positive (+), negative (−) and non-significant (0) and grouped by combination of GSE19743 and GSE182616 datasets. * P < 0.05, ** P < 0.01, *** P < 0.001.

**Supplemental Figure S3. COVID-19 single-cell RNA sequencing prolife.**

(A) Dotplot of cell marker genes in COVID-19 scRNA-seq. The dot color and size represents average expression and expressed percentage of each cell type.

(B) Cell proportion of each cell type among different disease statuses on sample-collection time. The error bar represented the 95% confidence interval and the y axis represented the cell percentage.

**Supplemental Figure S4. Mono-DC wing was the major cell source of SRC genes.**

(A) Boxplot of SRC genes score among each cell types in healthy subjects.

(B-D) DEGs GO enrichment of SRC+/-CD1C-CD141-DCs (B), cDCs (C) and monocytes (D).

(E) MHC-II Antigen presentation score of representative SRC+ Mono-DCs. SRC+ monocytes in asymptomatic subjects were compared to the other sub SRC+ cells. Wilcoxon test was applied and p value was adjusted by FDR.

(F) SRC genes expression dot plot in SRC+ Mono-DCs across disease statuses. The dot color and size indicated the scaled expression level and percentage.

(G) Boxplot of SRC genes score across disease statuses in CD1C-CD141-DCs, monocytes and cDCs. * P < 0.05, ** P < 0.01, *** P < 0.001, **** P < 0.0001.

**Supplemental Figure S5. CD1C-CD141-DCs sub cluster of critical COVID-19 patients is IFN pathway inactivated.**

(A) Barplot of top 24 TF regulons of SRC genes. The color indicated the max NES of TF.

(B) Network of top 24 TF and their target genes. The node color was annotated by TF and target genes. The node size represented the degree of each node.

(C) Heatmap of predicted STAT1 and IRF8 regulon activity and targeted gene expression of sub clusters of CD1C-CD141-DCs. The antigen-presentation- and IFN- related genes were marked on the left side.

(D) Representative GO enrichment barplot of STAT1 and IRF8 targeted genes.

(E) Boxplot of SRC (left), IFN alpha/beta (middle) and gamma (right) signaling score of CD1C-CD141-DCs across the different statuses. The critical COVID-19 was employed as the referent group. * P < 0.05, ** P < 0.01, *** P < 0.001, **** P < 0.0001.

**Supplemental Figure S6. CD1C-CD141-DCs sub cluster of critical COVID-19 patients is immune quiescent with other cells.**

(A) Dotplot of representative DEGs of IFN- and TNF-pathway related genes across sub clusters of CD1C-CD141-DCs. The dot color and size indicated the average expression level and expressed percentage of the relative cluster.

(B) Boxplot of CD4 expression across disease statuses in SC-RNA sequencing. Critical subjects were compared to the other subjects. P values were adjusted by FDR.

(C-F) Representative dot plot of DN DC incoming signaling, including TNF (C), ADGRE5-CD55 (D), ANXA1-FPR2 (E) and THBS1-CD36 (F). The dot size indicated the p value and the color represent communication probability. DN DC represents different sub clusters of CD1C-CD141-DCs. DN DC represents CD1C-CD141-DC. * P < 0.05, ** P < 0.01, *** P < 0.001, **** P < 0.0001.

